# Rapid monitoring for ecological persistence

**DOI:** 10.1101/2022.07.02.498308

**Authors:** Chuliang Song, Benno I. Simmons, Marie-Josée Fortin, Andrew Gonzalez, Christopher N. Kaiser-Bunbury, Serguei Saavedra

**Affiliations:** Department of Biology, Quebec Centre for Biodiversity Science, McGill University, Montreal, Quebec H3A 1B1, Canada; Department of Ecology and Evolutionary Biology, University of Toronto, 25 Willcocks Street, Toronto, Ontario M5S 3B2, Canada; Centre for Ecology and Conservation, College of Life and Environmental Sciences, University of Exeter, Cornwall Campus, Penryn TR10 9FE, UK; Department of Civil and Environmental Engineering, MIT, 77 Massachusetts Av., Cambridge, MA 02139, USA

## Abstract

Effective conservation of ecological communities requires accurate and up-to-date information about whether species are persisting or declining to extinction. The persistence of ecological communities is largely supported by its structured architecture of species interactions, known as an ecological network. While the persistence of the network supporting the whole community is the most relevant scale for conservation, in practice, we can only monitor small subsets of these networks due to logistical sampling constraints. There is therefore an urgent need to establish links between the small snapshots of data conservationists are able to collect, and the ‘big picture’ conclusions about ecosystem health demanded by policy makers, scientists and societies. Here we show that the persistence of small subnetworks in isolation — that is, their persistence when considered separately from the larger network of which they are a part — is a reliable probabilistic indicator of the persistence of the network as a whole. Our results are general across both antagonistic and mutualistic interaction networks. Empirically, we show that our theoretical predictions are supported by data on invaded networks in restored and unrestored areas, even in the presence of environmental variability. Our work suggests that coordinated action to aggregate information from incomplete sampling can provide a means to rapidly assess the persistence of entire ecological networks and the expected success of restoration strategies. This could significantly improve our ability to monitor progress towards achieving policy targets, such as those enshrined in the UN Convention on Biological Diversity.

## Introduction

To assess progress towards local, national, and international biodiversity conservation targets, it is crucial to understand how ecological communities are being impacted by global change and how well they are responding to conservation interventions^1,2^. Biomonitoring—the process of regularly measuring ecosystems to track changes in different indicators over time—therefore has a critical role to play in guiding conservation policy and practice.

Perhaps, the most fundamental indicator of the health of an ecological community is whether any of its component species are declining to extinction^3^. Thus, for conservation scientists and decision makers, community persistence—the probability that a community can sustain positive abundances of all its constituent populations over time—is one of the most important properties to know about a community ^4,5^.

One widespread approach to measure community persistence relies on species interaction networks, where nodes, representing species, are joined by links, representing biotic interactions, such as predation or pollination^6^. These networks are then used as a ‘skeleton’ for simulating population dynamics, from which persistence is then measured. While this approach is powerful, collecting interaction network data using traditional field methods is expensive and time consuming, often requiring huge investments to approach sampling completeness^7–9^. For example, studies have shown that even with exhaustive sampling effort that covered 80% of pollinator fauna, it was only possible to capture 55% of the interactions in a plant-pollinator network and it was estimated that sampling effort would need to increase by five times to record 90% of the interactions ^10^. Similarly, while DNA barcoding and metabarcoding approaches offer the prospect of achieving complete sampling of networks more easily, at present, these methods are far from the norm ^11^, requiring significant technical expertise, sampling protocols and infrastructure with costs that can be comparable to traditional field approaches^11^.

The expense and time involved in collecting community-level data, such as sampling interaction networks, is in direct contrast to the needs of biomonitoring and the frugal realities of conservation practice ^7^. Biomonitoring relies on indicators that are sufficiently quick and cheap to sample that regular ‘snapshots’ of ecosystem status can be taken. Thus, although persistence is a fundamental property of communities that we wish to monitor, time and cost constraints mean that sampling the required interaction network data with sufficient temporal, spatial, and taxonomic resolution is impractical in all but the most well-resourced contexts.

It has been suggested that interaction networks can be monitored cost efficiently if only a small subset of interactions – a subnetwork – is sampled^7,12^. However, it is not known whether there is any consistent relationship between the persistence of a subnetwork and the persistence of the larger network that the subnetwork comes from. If such a relationship could be established, then whole-network persistence could be inferred from subnetworks that require much less effort to sample. In turn, the prospect of rapid and affordable biomonitoring of network persistence would become feasible.

Here, we provide a solution to diagnosing the persistence of ecological networks without the need for extensive sampling. Specifically, we show that the persistence of a network as a whole is linked to the persistence of small-scale subnetwork structures that require much less sampling effort to monitor (Figure 1). We use coexistence theory to show that mutualistic and antagonistic interaction networks that are persistent as a whole are comprised of subnetworks that would persist in isolation. That is, they are composed of subnetworks that would persist even if they were not embedded within the larger network of interactions. We term these subnetworks *persistent in isolation*. We show that reliable conclusions about the persistence or non-persistence of the whole network can be made from only sampling 2-4 subnetworks. We corroborate our results using empirical data on invaded mutualistic interaction networks ^13^. We show that persistent ecological networks, despite their spatio-temporal variability, contain more persistent subnetworks in isolation in areas where invasive species have been removed through restoration action than in areas where there has been no restoration. Finally, we discuss our results in light of new opportunities for rapid biomonitoring of network persistence.

**Figure 1:**
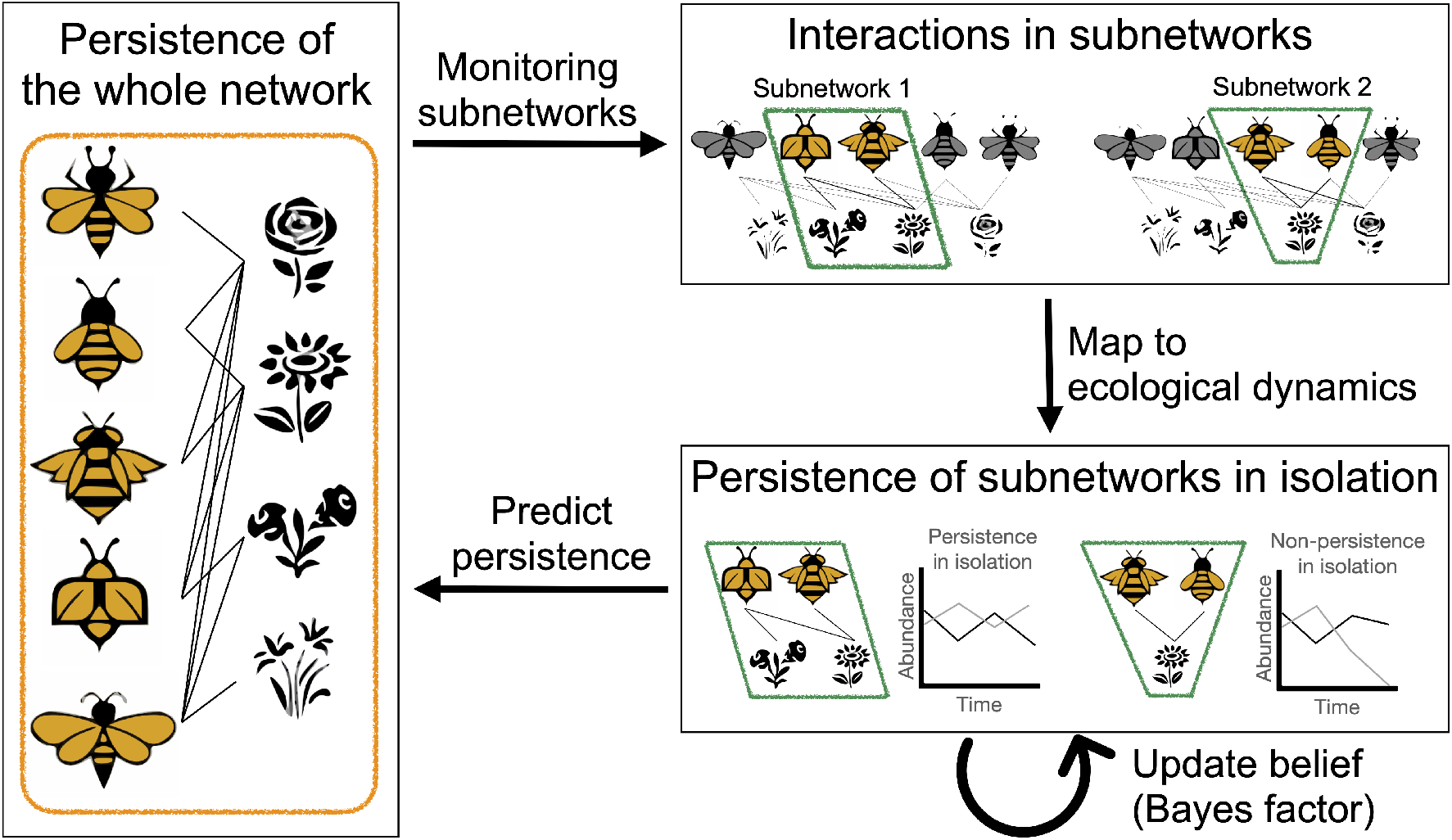
Illustrated scheme to detect network persistence from monitoring subnetworks. For illustrative purposes, we consider a hypothetical network structure of a mutualistic network consisting of five pollinator species and four plant species (orange box). Persistence is a network-level property, which makes the network scale the most relevant scale for measuring persistence. However, time and cost constraints and sampling biases limit our knowledge of the network structure. Thus, we most often observe small-scale subnetworks called motifs (two green boxes) that are embedded into the larger network. ‘Embedded persistence’ refers to persistence of these motifs that are part of larger networks. However, given that the whole network is unobservable, we can instead study whether the subnetworks can persist in isolation, removed from the wider network context, which we call the ‘persistence in isolation’ of a subnetwork. While it is clear that the persistence of the network determines the embedded persistence of subnetworks, here we show that the persistence in isolation of subnetworks is linked to the persistence of the network as a whole, and we can update our belief on the persistence of the whole network from observing the persistence in isolation of the subnetworks.

### Linking network and subnetwork persistence

The challenge we face is knowing whether the persistence of a large species interaction network can be inferred by only observing the structure of one or more subnetworks (also known as interaction motifs). The structural approach in ecology^14,15^ provides a theoretical solution to this problem by connecting the persistence of the network as a whole with that of its subnetworks. The central concept in the structural approach is the *coexistence domain* (*D*)—the full range of conditions (parameter space) under which all species in a network can coexist, given a particular network structure^16^. Hence, the persistence of a network as a whole can be studied using the coexistence domain of its underlying network structure (*D*_whole_). Similarly, to understand the persistence of a subnetwork, we can study the coexistence domain of its underlying subnetwork. This can be done in two ways, as illustrated in Figures 1 and S1. First, we can study the subnetwork as part of the larger network in which it is embedded; formally, this is the projection of the coexistence domain of the network onto the subnetwork 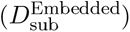 (Figures 2 and S1). Second, we can study the coexistence domain of the subnetwork in isolation, removed from the larger network of which it is a part 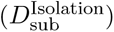 (Figures 2 and S1).

**Figure 2:**
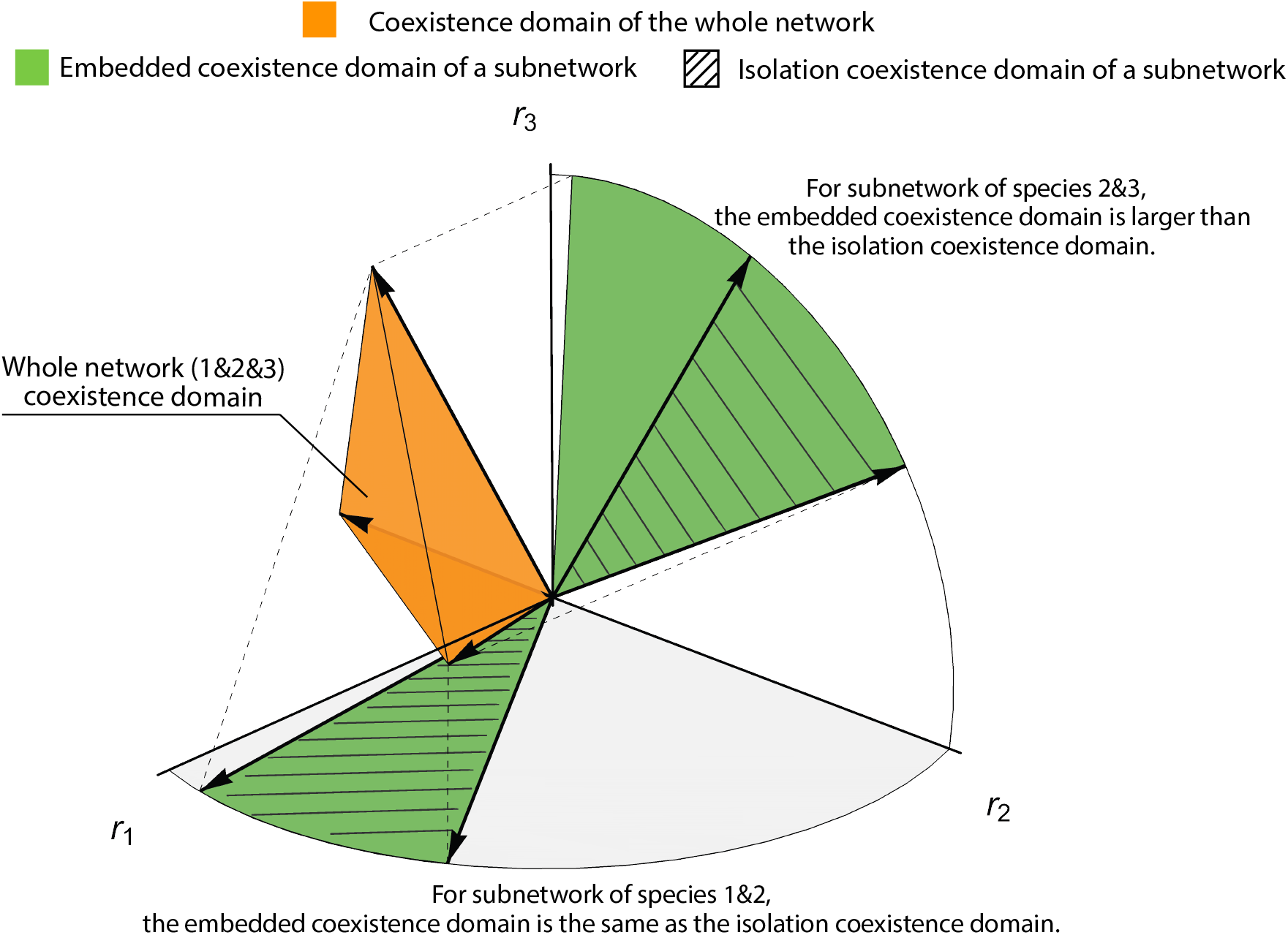
A structural approach to link persistence across scales. We establish the link between the network and the subnetwork scales via the concept of the *coexistence domain*: the region of all parameters in which a given set of species coexists. For illustrative purposes, we consider a hypothetical 3-species network. If we want to understand persistence of the network as a whole, we should study the coexistence domain of the network where all species coexist (orange region). If we want to understand the isolation persistence of a subnetwork, we should study the coexistence domain of the subnetwork in isolation i.e., removed from the larger network of interactions in which it is embedded (hashed region). If we then want to understand the persistence of the network as a whole, but we can only observe a subnetwork (embedded persistence), then the observed coexistence domain for the subnetwork is the projection of the network’s coexistence domain into the subnetwork’s (light green region). The projection of the network’s coexistence domain is always larger than (e.g., subnetwork with species 2 and 3) or equal to (e.g., subnetwork with species 1 and 2) the subnetwork’s coexistence domain.

In field surveys, where only a subset of a network can be sampled, the network in which that subset is embedded is not observed—only 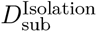 is known. Thus, the question is whether the true unobservable coexistence domain of the subnetwork 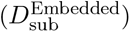 is equivalent to the observed coexistence domain of the subnetwork in isolation 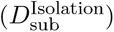 and whether 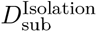 can be used to infer the persistence of an unobservable network as a whole (*D*_whole_).

We formalize 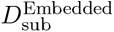 and 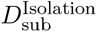 using the structural approach. The structural approach applies to any population dynamics that are topologically equivalent to Lotka-Volterra dynamics ^5,17^. Note that any quasi-polynomial dynamics, which include a large class of ecological models, can be equivalently mapped into Lotka-Volterra dynamics ^18^. Here, we assume that a network has *S*-interacting species in total and is governed by the Lotka-Volterra dynamics^19^:

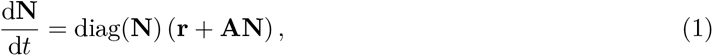

where **N** is the vector of species densities and **A** is the matrix of species interactions. Under the governing population dynamics, the coexistence domain of a network is given by^16^:

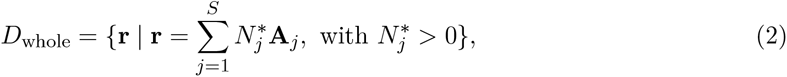

where **A**_*j*_ denotes the *j*-th column of the interaction matrix **A**. The coexistence domain of a subnetwork with the set of species 𝒫 in isolation is given by

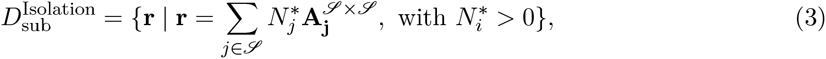

where **A**^𝒫×𝒫^ is the sub-matrix of **A** that only has species in the set of species 𝒫. In turn, the coexistence domain of a subnetwork with the set of species 𝒫 when embedded in the larger network is given by

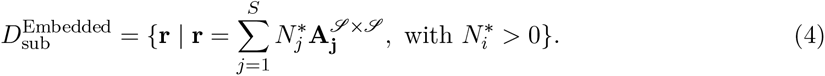

It is important to note that 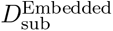 (Eqn. 4) is always greater than or equal to 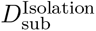 (Eqn. 3).This is because the summation in Eqn. 4 is done across all columns, while the summation in Eqn. 3 is only done across a subset of all the columns. For example, in Figure 2, 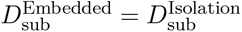 for the subnetwork with species 1 and 2, while 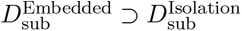 for the subnetwork with species 2 and 3. This result implies that observing only a subset of species without its wider network context (i.e., 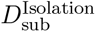) never overestimates the true coexistence potential of this subset when embedded into the larger network of interactions (i.e., 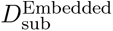). This result is general and does not depend on types of species interactions or the number of species.

### Monitoring network persistence from subnetwork persistence

To study the relationship between subnetwork persistence in isolation and network persistence as a whole, we borrow concepts from statistical physics and test whether a macroscopic (whole-network) phase transition is reflected in drastic changes of mesoscopic (subnetwork) properties ^20^. At the macroscopic scale, the network as a whole only exhibits two phases: persistence or non-persistence. At the mesoscopic scale, a subnetwork has an additional key property: whether the persistence of the subnetwork is identical in isolation and when embedded in the larger network. An embedded subnetwork is persistent if, and only if, the network as a whole is persistent. Thus, if the persistence of a subnetwork is identical in isolation and when embedded in the larger network, then the subnetwork’s persistence (or not) in isolation also indicates whether the network as a whole persists (or not). This property is always true for the subnetworks with 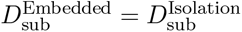, and is true with probability 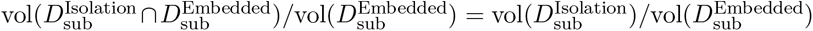 for all subnetworks. This relationship between 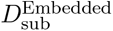 and 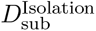 suggests the possibility that the persistence of *one* subnetwork can be a *probabilistic* indicator of the persistence of the network as a whole. Furthermore, the persistence of *many* subnetworks can be an almost-*deterministic* indicator of the persistence of the network as a whole.

From the above, we hypothesize that the proportion of subnetworks that are persistent in isolation is different between persistent and non-persistent networks. To formally test this hypothesis, we simulate theoretical networks, and analyze the persistence in isolation of their constituent subnetworks as the network is moved along a gradient from persistence to non-persistence. For brevity, we focus on mutualistic interaction networks in the main text, but our approach can equally be applied to antagonistic interaction networks, such as predation and parasitism (see Appendix B)^6,21^. First, we generate theoretical mutualistic networks with 10 pollinators and 8 plants (see Appendix B for the robustness of our results to different network sizes). For illustrative purposes, we present here Erdős–Rényi structures, however, our results apply and are consistent for other network structures, such as nested networks (Appendix B). We then follow an ecologically-motivated parameterization that has been widely adopted in the literature^22–25^. The interaction matrix of a mutualistic network can then be partitioned as

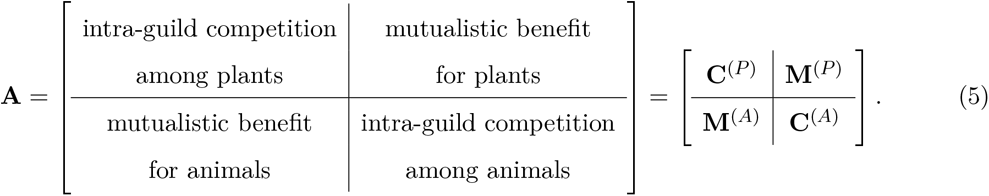

The mutualistic benefit (**M**^(*P*)^ and **M**^(*A*)^) between species *i* and *j* is parameterized as 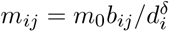 if species *i* and *j* directly interact and *b*_*ij*_ = 0 otherwise, *m*_0_ represents the overall strength of mutualistic interaction, *d*_*i*_ represents the number of interaction partners, and *δ* represents the mutualistic trade-off. The intra-guild competition (**C**^(*P*)^ and **C**^(*A*)^) between species is parameterized using a mean-field approach where we set the intraspecific competition 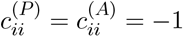 and interspecific competition 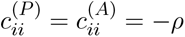.

We systematically sample inside and outside the coexistence domain of simulated networks (i.e., across two macroscopic phases: persistence and non-persistence) and check the persistence in isolation of all constituent subnetworks at each sampling point (Figure 3A). We adopt the normalized distance of the sampled point from the centroid of the coexistence domain as the tuning parameter of the phase transition. Specifically, the centroid has a normalized distance 0, all points on the border of the coexistence domain have normalized distance 0.5, and all points inside (resp. outside) have normalized distances less (resp. greater) than 0.5 (Figure 3A).

**Figure 3:**
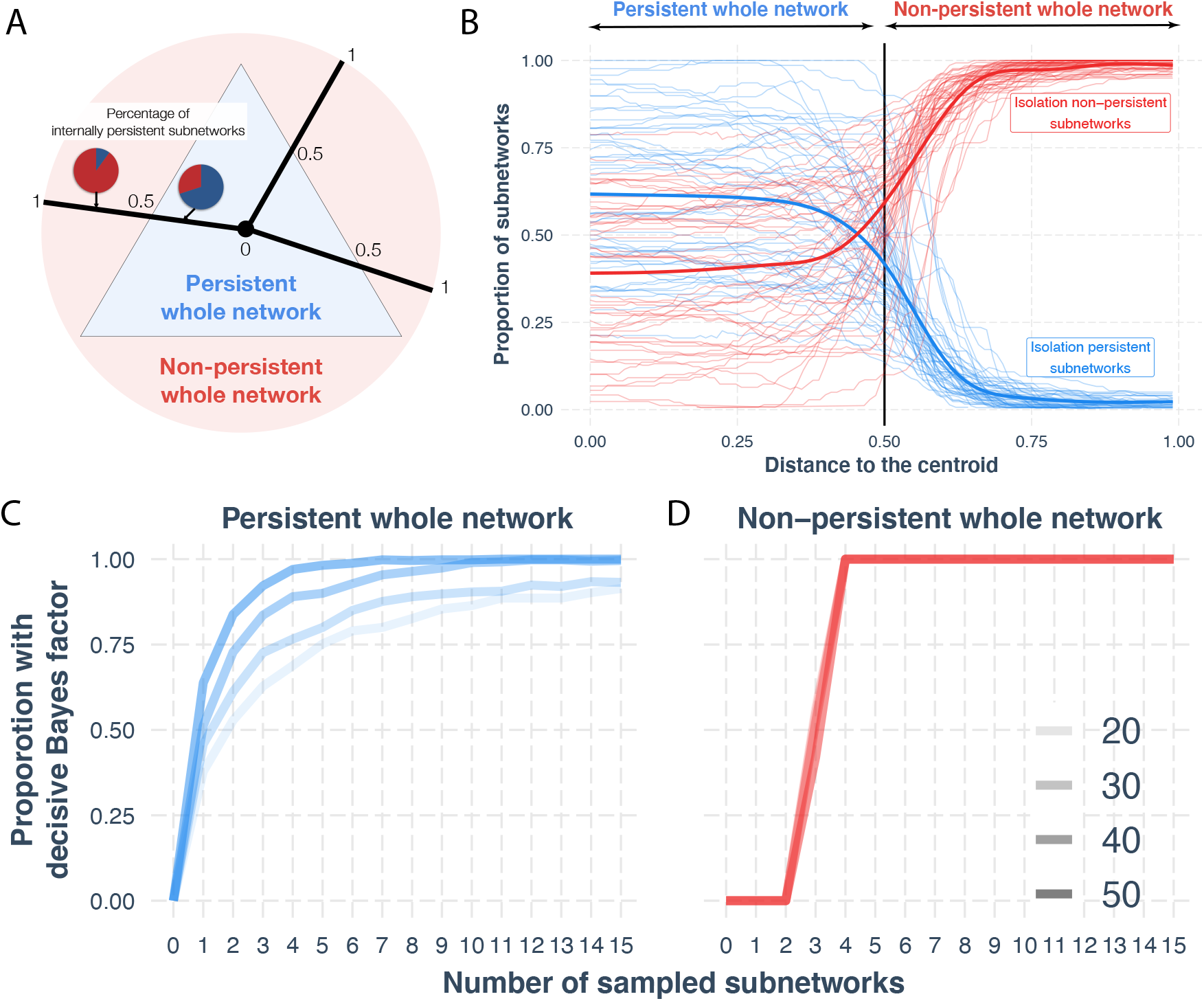
Generic phase transition of persistence across scales. (**A**) Schematic illustration of the structure of our experiment. For a given network, we measure the proportion of subnetworks (motifs) that are persistent along ‘transects’ that span inside and outside the coexistence domain of the network. Within the coexistence domain, the network is persistent; outside the coexistence domain the network is non-persistent. The transects span a tuning parameter that ranges from 0 to 1, with 0 being at the centre of the coexistence domain and 0.5 being at the boundary of the coexistence domain. (**B**) Here, we explore the subnetwork mechanisms driving network persistence as a whole. The *x* axis denotes a tuning parameter moving the network from non-coexistent (non-persistent) to coexistent (persistent). The *y* axis denotes the proportion of subnetworks (motifs) in the network that persist (blue) and do not persist (red) in isolation. The whole network is persistent in the left half and non-persistent in the right half. Each thin line represents one simulation along a ‘transect’, as shown by the black line in panel A (50 are shown here), and the think lines denote the average. We see a transition in the proportions of persistent and non-persistent subnetworks as the network transitions from persistence into non-persistence. This transition shows that a persistent network is primarily composed of persistent subnetworks, and *vice versa*. This phase transition is generic in nearly almost all simulations (see Appendix B). (**C**)-(**D**) Monitoring network persistence as a whole from subnetworks. Panel (C) shows the case for persistent network (left side of Panel B), while Panel (D) for non-persistent case (right side of Panel B). The *x* axis shows the number of randomly sampled subnetworks (from 3 to 5 species). The *y* axis shows the proportion of samples that reaches decisive Bayes factor that supports that the network is persistent (in Panel C) or non-persistent (in Panel D). The transparency of the lines denotes different network sizes (from 20 to 50). We find that it is generally easier to statistically determine if the network as a whole is not persistent from monitoring subnetworks.

We characterize subnetworks using bipartite motifs ^26,27^. Bipartite motifs are small subnetworks containing between two and six species, with all species having at least one link^26,27^. Bipartite motifs describe all possible unique subnetwork topologies up to six species. As described above, we enumerate all the motifs present in each network along the gradient of persistent and non-persistent networks (Figure 3A) and calculate both the persistence in isolation and embedded persistence of each motif. Although the number of motifs scales exponentially with network size, we expect to see a smooth, rather than sharp, phase transition in the persistence of motifs along the gradient, given the small network size in our simulations^20^.

Corroborating our hypothesis, as networks were moved along a gradient from network persistence to non-persistence (Figure 3A), we identified a phase transition of persistence across scales of networks and subnetworks (Figure 3B). That is, when a network as a whole is persistent, the majority of its constituent subnetworks (motifs) must themselves be persistent in isolation. Conversely, when the network as a whole is not persistent, the majority of its constituent subnetworks are not persistent in isolation. Overall, we found that, when 2-4 of a set of randomly sampled subnetworks are persistent, the network as a whole is persistent. Simulations suggest that this result is general across mutualistic and antagonistic interaction networks, and networks of different sizes (Appendices B). We also provide a heuristic explanation of the emergency of the phase transition (Appendix C).

These theoretical findings additionally corroborate that the presence of a *single* persistent subnetwork in isolation is a probabilistic indicator of the persistence of the network as a whole, while the presence of *many* persistent subnetworks in isolation is a nearly deterministic indicator (with exponentially decreasingly error) of the persistence of the network. To formalize this idea, we provide a Bayesian approach to update our beliefs about the persistence of the network from whether subnetworks are persistent in isolation. Specifically, we compute the Bayes factor: the ratio of the posterior likelihood of the network being persistent as a whole over being non-persistent given the observed persistence in isolation of sampled subnetworks. A Bayes factor larger than 10^2^ can be considered decisive evidence that the network is persistent, while a Bayes factor smaller than 10^−2^ can be considered decisive evidence that the network is non-persistent^28^. Figure 3C-D shows how many subnetworks (containing 3-5 species) we need to sample to reach decisive evidence about whole-network persistence (more details on the simulations in Appendix D). When the network as a whole is persistent (Figure 3C), the number of required sampled subnetworks decreases with increasing network size (per statistical laws as roughly only half of subnetworks are persistent in isolation). In contrast, when the network is not persistent, the number of required sampled subnetworks is much lower and does not vary with network size (as most subnetworks are not persistent in isolation). Overall, the results in Figure 3C-D show that only 2-4 subnetworks are needed to be reasonably confident about the persistence or non-persistence of a whole network.

### Empirical analysis

While it is difficult to directly empirically test the phase transition resulting from the theoretical analysis above, a testable prediction from our result is that a network as a whole is more likely to persist if it contains more persistent in isolation subnetworks. To test this, we analyzed a temporal dataset of 8 plant–pollinator networks from the Seychelles, each sampled over 8 consecutive months (144 pollinator species and 38 plant species in total). In this dataset, half of the networks are subject to disturbance in the form of invasive plants, while the other half have had invasive plant species removed through restoration action ^13^. We expect restored sites to have higher persistence than unrestored invaded networks and, as a consequence, we expect these restored networks to contain more persistent in isolation subnetworks (motifs) than invaded networks.

We measure the persistence in isolation of subnetworks as the size of their coexistence domains. Note that comparisons between subnetworks can easily be confounded by different parameterizations. That is, small changes in parameters used to calculate the coexistence of different subnetworks can significantly influence conclusions^29,30^. To ensure a fair comparison, we focus on subnetwork pairs that are the transpose of each other, and parameterize both subnetworks in the pair identically (Figure S9). For example, we compare the subnetwork with two top nodes connected to a single bottom node, with the subnetwork containing two bottom nodes connected to a single top node. In ecological terms, this might be comparing the subnetwork describing two pollinators visiting one plant to the subnetwork with one pollinator visiting two plants. In total, we have seven pairs of subnetworks that range from 3 species to 5 species each, and 10 pairs of subnetworks with 6 species each (Figure S9). Note that despite the interaction matrices for each pair of subnetworks being the transpose of each other, the coexistence domains of the two subnetworks are different because of, and only because of, their trophic constraints^16,31^. In other words, plants always have positive intrinsic growth rates in our bipartite networks.

For each different subnetwork, we compute the size of its coexistence domain by systematically exploring the parameter space, defined by three parameters: the mutualistic trade-off *δ*, the overall strength of mutualistic interactions *m*_0_, and the intra-guild competition strength *ρ*. We then examined the empirical frequency of each subnetwork (motif) in the observed mutualistic networks. We first computed the raw number of subnetworks present in each empirical mutualistic network using the *bmotif* R package ^27^. Next, we compute the *z*-score of the empirical subnetwork frequencies relative to the Erdős–Rényi null model ^32,33^. Next, we compare the patterns of these null-corrected subnetwork frequencies between unrestored (invasive species present) and restored (invasive species removed) sites ^13^. Lastly, we investigate if these patterns remain if we can only monitor a subset of all species; this tests whether conclusions about whole-network persistence can be made from easy-to-sample subnetworks.

We expect that networks, where invasive species had been removed, would be more persistent (i.e., with a larger coexistence domain) and thus would have higher numbers of persistent in isolation subnetworks. Figure 4A shows that the restored (undisturbed) networks have a higher over- representation of persistent in isolation subnetworks (motifs) compared to the unrestored (disturbed) networks. This pattern is robust through the whole sampling period, despite the reorganization of species interactions in the network. Importantly, Figure 4B shows that these patterns remain qualitatively true even if we can only monitor the interactions of a small subset of all locally present species. This shows that we can gain insights about the network as a whole using only information from a subset of that network that is much quicker to sample.

**Figure 4:**
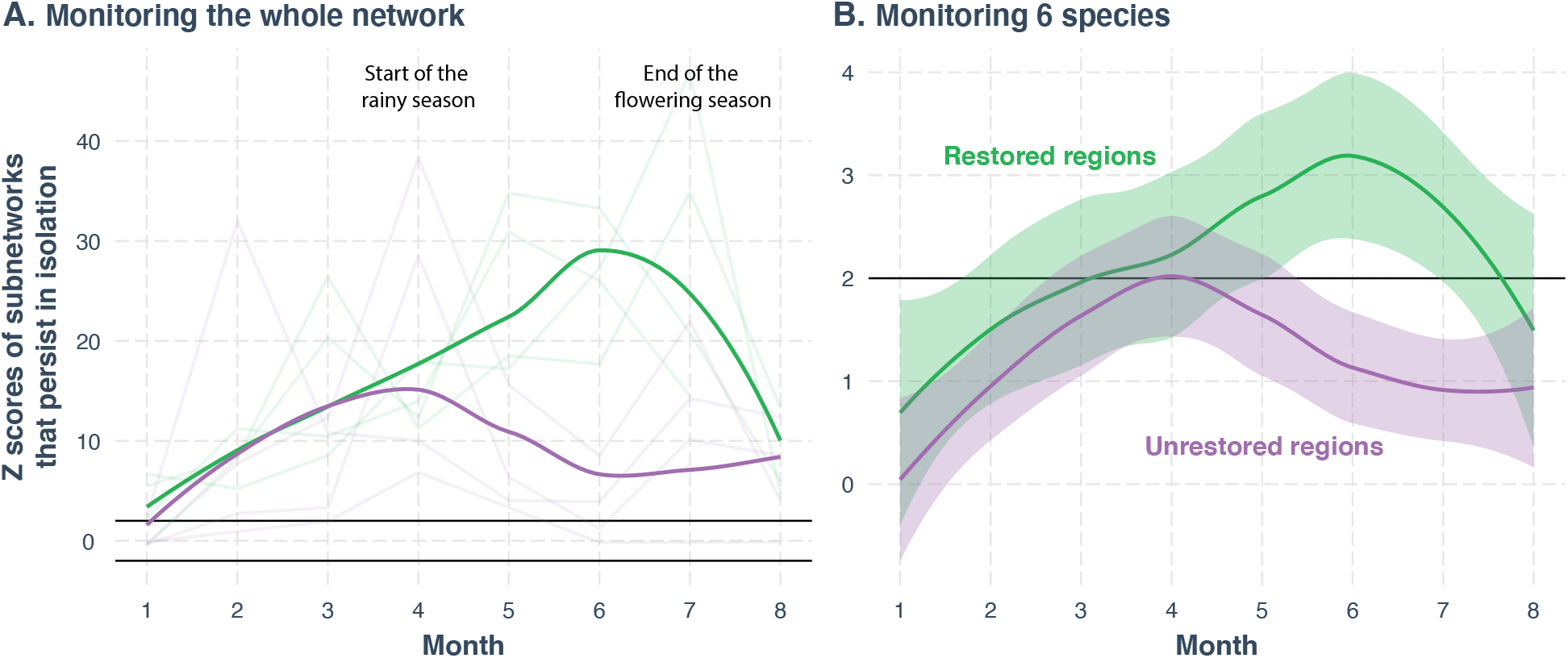
Restored mutualistic networks contain more subnetworks with higher likelihood of persistence in isolation than unrestored disturbed networks. We use a temporal dataset with 64 networks (8 networks sampled over 8 months) located on the granitic island of Mahé, Seychelles ^13^. The *x* axis denotes the eight consecutive months between September 2012 and April 2013. The *y* denotes the *z*-scores of a given subnetwork (over- or under-representation of empirical motif frequency compared to motif frequency in randomized networks). The black horizontal line corresponds to the threshold above or below which a subnetwork (motif) occurs significantly more than random (z-score = 2). The green lines correspond to the restored networks, while the purple lines correspond to the disturbed and unrestored networks. (**A**) Monitoring the network. Each translucent light line corresponds to a different network. The thick lines correspond to the average across 4 different networks. The persistent in isolation subnetworks are significantly over-represented in the larger networks, and the over-representation is stronger in restored networks than in disturbed networks. These patterns are consistent with the natural history of these plant-pollinator networks. (**B**) Monitoring a subset of the network. Suppose that we cannot monitor the whole network but only a subset of it. Here we show the case for monitoring 6 species (see Appendix F for other numbers of monitored species). We find that all the qualitative patterns linking subnetworks and the network in Panel (A) remains. This shows that we can monitor a subnetwork (6 species of the whole network) and then study the persistence in isolation of its subnetworks (3-5 spices in the monitored 6 species), which would provide useful information of the network as a whole.

## Discussion

We have uncovered a generic probabilistic link across organizational scales of ecological networks: an ecological network is more likely to be persistent when a majority of its subnetworks are persistent in isolation, and *vice versa*. This establishes a theoretical foundation to rapidly monitor the persistence of communities by sampling only small numbers of interactions in subnetworks. Given cost and time constraints common to much conservation work, it is only practical to have high quality data on a small fraction of all locally present species and interactions. Our results have shown that determining the persistence in isolation of a few subnetworks can lead to robust conclusions about the persistence of the network as a whole: if 2-4 randomly sampled subnetworks are persistent, then the network as a whole is likely persistent. Our monitoring scheme is insensitive to the size of the whole network, which is in direct contrast to prohibitively expensive sampling of the whole network (estimated to be exponential efforts with the size of the whole network^10^). Our results help to address a fundamental challenge to make predictions under limited observations in conservation management^34,35^.

Besides the application to rapid biomonitoring, our work also connects two separate schools of thought on ecological networks defined by distinct ecological scales. One school has focused on the mesoscopic scale, providing a more mechanistic understanding through the study of specific small subnetworks or motifs (often called ‘trophic modules’), such as apparent competition^36–40^. In contrast, the other school has focused on the macroscopic scale, studying the interaction networks of entire networks to provide an ecologically more relevant link, but suffering from being coarse-grained and highly phenomenological ^23,41–43^. Unfortunately, these two schools have little cross-talk. The few theoretical studies on this topic suggest there is no deterministic, one-to-one link that maps results across the two schools ^44–46^. We confirm that such a deterministic link does not exist. However, we found that a generic probabilistic link does exist (Figures 2 and 3). This generic link provides an opportunity to take advantage of both schools. For example, we have taken a phenomenological approach with the trophic constraints to explain why some subnetworks are more persistent in isolation. However, a rich literature has cataloged and explained why some subnetworks are more persistent in isolation, ranging from sign stability^47,48^ to consumer-resource relationships^40,49^, and then applied it to explain observed patterns of species-rich ecological networks^39,50,51^. Our results have justified that such practice is probably approximately correct ^52^.

We have provided a Bayesian approach to update our belief on the persistence of ecological networks (Figure 1). Our belief is quantified by the Bayes factor—the ratio of the posterior probability of the monitored network being persistent over not being persistent. The Bayes factor is consistently updated by our knowledge of species’ life history (as priors) together with persistence-in-isolation criteria of the observed subnetworks (as evidence). For simplicity, we have assumed the prior Bayes factor to be 1 (i.e., equal likelihood for the monitored network to be persistent or not), but we can set more realistic prior Bayes factor in applied research by incorporating indigenous and expert knowledge ^53,54^. We have found that conclusions about whole network persistence or non-persistence can be made from only a handful of subnetworks (Figure 3).

Our key theoretical prediction—that subnetworks with a higher probability of persistence in isolation are over-represented in more persistent networks—is supported by empirical temporal mutualistic networks. These empirical results agree with the natural history details of these systems. For example, Figure 4 shows how restored networks have only slightly higher frequencies of persistent subnetworks than unrestored networks until month four, when the two treatments diverge, and persistent subnetworks substantially increase in number in restored sites relative to unrestored (disturbed) sites. Networks in the two treatments then converge at the end of the flowering season. This divergence in month four corresponds to the start of the rainy season in the Seychelles. Rains create a patchy distribution of resources. In the unrestored sites, free movement of pollinators between patches is hindered because invasive species grow densely, disrupting the ability of pollinators to for-age effectively^13^. If pollinators cannot find resources, or cannot move freely between individuals and species, this increases competition and reduces niche complementarity, which likely causes reduced persistence.

Overall, the match between theoretical predictions and empirical observations shows that information about the persistence of a species-rich ecological network is encoded in its small subnetworks and can be recovered through an appropriate theoretical lens and statistical analysis. This finding has significant application to biodiversity monitoring as indicators of ecological persistence and biodiversity intactness are key foci of monitoring networks around the world. The method we have provided here has the potential to save time and effort and accelerate our ability to detect and mitigate unwanted change to the structure and function of ecological networks.

## Acknowledgements and funding sources

B.I.S. is supported by a Royal Commission for the Exhibition of 1851 Research Fellowship. A. G. is supported by the Liber Ero Chair in Biodiversity Conservation and the Quebec Centre for Biodiversity Science. M.-J. F. thanks the Natural Sciences and Engineering Research Council of Canada and a Canada Research Chair for their support. Funding to S.S. was provided by NSF grant No. DEB-2024349.

